# Humoral Immune Dysregulation Defines High-Risk Cirrhotic Ascites

**DOI:** 10.64898/2026.09.25.754574

**Authors:** Arielle Klepper, Matt S. Zinter, Tamara Roach, Sukhman Sidhu, Zoe Berman, Ishaan Dureja, Ravi Dandekar, Sammer Ganem, Karina Perlaza, Bryan Castillo-Rojas, Kelsey Zorn, Jacquelyn J. Maher, Jody Baron, Matthew Yocum, Andrew R. Lai, Jennifer Lai, Joseph DeRisi

## Abstract

Fluid accumulates within serosal cavities in diverse disease states, where it creates biologically active and potentially pathologic niches. How these expanded microenvironments reflect and shape disease biology remains incompletely resolved. Here, we investigated human ascites fluid (*n* = 118), a hallmark of decompensated cirrhosis, to determine whether signatures of microbial exposure and local host response distinguish outcomes. Cirrhotic ascites was characterized by barrier remodeling, depletion of plasma cell-associated programs, and enrichment of oropharyngeal microbes. Surprisingly, detection of oropharyngeal microbes in ascites was not associated with adverse outcomes. Instead, the ascites of cirrhotic patients with poor outcomes was marked by profound plasma cell depletion and significantly reduced dimeric IgA, despite the presence of antigen-experienced B cells. Together, these findings identify local B cell dysregulation as a previously underappreciated component of cirrhosis-associated immune dysfunction. Impaired local humoral immunity, rather than microbial translocation alone, emerges as a feature of high-risk decompensated cirrhosis.

## Introduction

The movement of fluid between the vascular and extravascular spaces was mechanistically described in Ernest Starling’s seminal work in 1896^1^, and disruption of these forces can result in pathologic fluid accumulation across diverse tissue and serosal compartments^2^. The hemodynamic forces underlying fluid accumulation, such as increased hydrostatic pressure associated with portal hypertension, are well established. In contrast, the molecular and cellular composition of these expanded fluid compartments has remained comparatively understudied. Emerging evidence demonstrates that these fluids are not merely passive consequences of disease, but instead represent biologically active niches characterized by disease-specific cellular, microbial, and immune processes, some of which have been associated with treatment response and clinical outcomes^3–6^.

Expansion of fluid within the peritoneal cavity, termed ascites, has been recognized since antiquity^7^, and occurs across a broad range of human diseases, including cirrhosis, malignancy, infection, and cardiometabolic disorders. Cirrhosis accounts for 80–85% of all ascites^8^. Despite this predominance, studies of cirrhotic ascites have lagged far behind those characterizing the molecular landscape of ascites due to malignancy^4,9–11^. This knowledge gap is particularly consequential because the development of ascites marks a major transition from compensated to decompensated cirrhosis and is accompanied by a substantial deterioration in prognosis^12^.

Current models of acute decompensation in cirrhosis point to portal hypertension – elevated portal venous pressure caused by increased resistance to flow through the cirrhotic liver – together with microbial translocation, infection, and immune dysregulation as interconnected processes associated with disease progression and adverse outcomes^13,14^. However, the molecular features that distinguish clinical trajectories among patients with decompensated cirrhosis are not fully understood.

Spontaneous bacterial peritonitis (SBP), an infection of ascitic fluid in the absence of an identifiable intra-abdominal source, directly implicates the ascites compartment as a site of clinically consequential host–microbe interactions in cirrhosis.^15^ Foundational studies in animal models established that translocation of gut microbes into ascites contributes directly to SBP, providing a mechanistic link between barrier dysfunction, microbial exposure, and local host immunity.^16–18^ Beyond infection, foundational studies in mice have established the peritoneal cavity as a distinct immunologic niche, home to specialized populations of resident macrophages and B cells^19,20^, which contribute to local immune defense and natural antibody production^21,22^. Studies of oncologic ascites suggest that this compartmentalization extends to humans, revealing immune populations, transcriptional states, and antigen-receptor repertoires distinct from those in peripheral blood^23^. In patients with cirrhosis, human counterparts of murine resident peritoneal macrophages have been identified, with macrophage activation linked to outcomes in SBP^24^. Despite the close relationship between cirrhosis, ascites, and infection, the broader host–microbe biology in cirrhotic ascites remains incompletely resolved.

Meta-transcriptomic profiling allows for an unbiased survey of ascites, coupling host response and microbial detection for an integrated read-out of host/microbe dynamics. This approach is particularly well suited for delineating which elements of the host-microbe interaction are associated with poor outcomes. Here we apply meta-transcriptomics to deliver an integrated molecular landscape of ascites and identify unique features of cirrhotic ascites – increased barrier remodeling, decreased plasma cell signatures, together with an increase in oral cavity associated microbes. However, microbial burden did not distinguish patients with adverse outcomes. Through deep orthogonal validation, we confirmed marked plasma cell depletion in cirrhotic ascites and linked impaired local humoral immunity to adverse outcomes. Reduced dimeric IgA distinguished high-risk disease and was associated with death or liver transplantation within 90 days.

## Results

### Cirrhosis leads to remodeling of the ascites microenvironment

To define the molecular and cellular features of cirrhotic ascites, we studied 118 hospitalized patients with ascites, including 65 with cirrhotic and 53 with non-cirrhotic ascites. Our investigation proceeded in three phases (Figure 1). In an initial discovery phase, we performed transcriptomic profiling of 10 mL of ultracentrifuged ascites from a subgroup of patients (*n* = 30) to identify candidate signatures distinguishing cirrhotic from non-cirrhotic ascites. We next validated the identified signatures in a separate group of patients from the same prospective cohort (*n* = 88), assessing both transcript and protein levels from 1 mL of unprocessed ascites. In the third phase, we performed deep immune profiling of cells from 1 liter of fresh ascites isolated from a nested subset of patients (*n* = 8). Clinical characteristics were incorporated throughout the analyses (Table 1a), while prospectively collected 90-day outcomes were used in subsequent mortality analyses (Figures 3 and 5; Table 1b).

**Figure 1.**
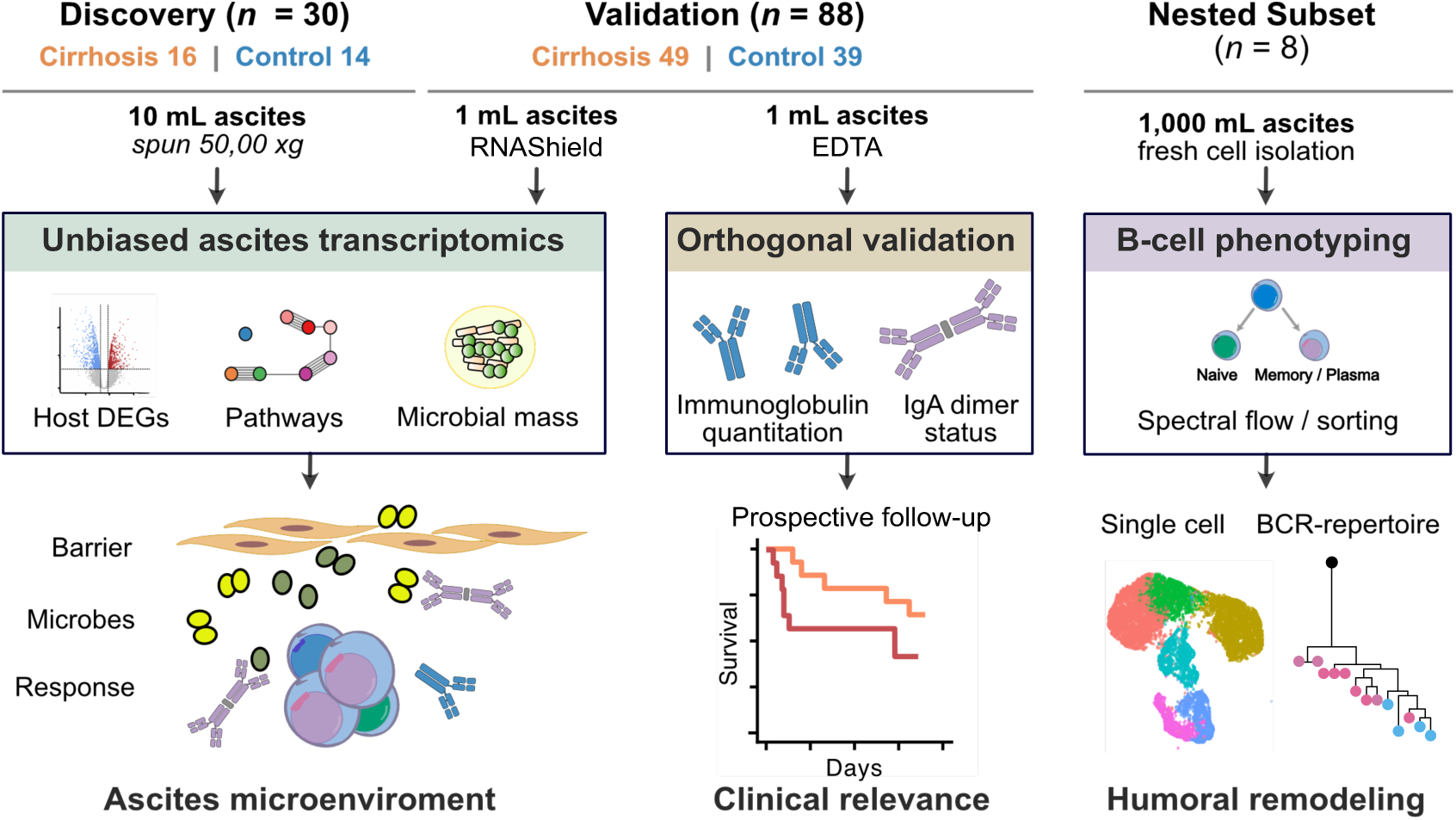
Study design and profiling of the ascites microenvironment. The ACES cohort included a discovery cohort (n = 30; 16 cirrhosis, 14 non-cirrhosis), an independent validation cohort (n = 88; 49 cirrhosis, 39 non-cirrhosis), and a nested deep-phenotyping cohort using 1 L of fresh ascites (n = 8; 4 cirrhosis, 4 non-cirrhosis). Discovery and validation studies integrated meta-transcriptomic host-gene and microbial profiling with orthogonal immunoglobulin measurements. The nested cohort underwent spectral flow cytometry and B-cell sorting, with single-cell RNA and paired B-cell receptor (BCR) sequencing in four patients. Molecular and cellular features were integrated with prospectively collected clinical outcomes.

**Figure 2.**
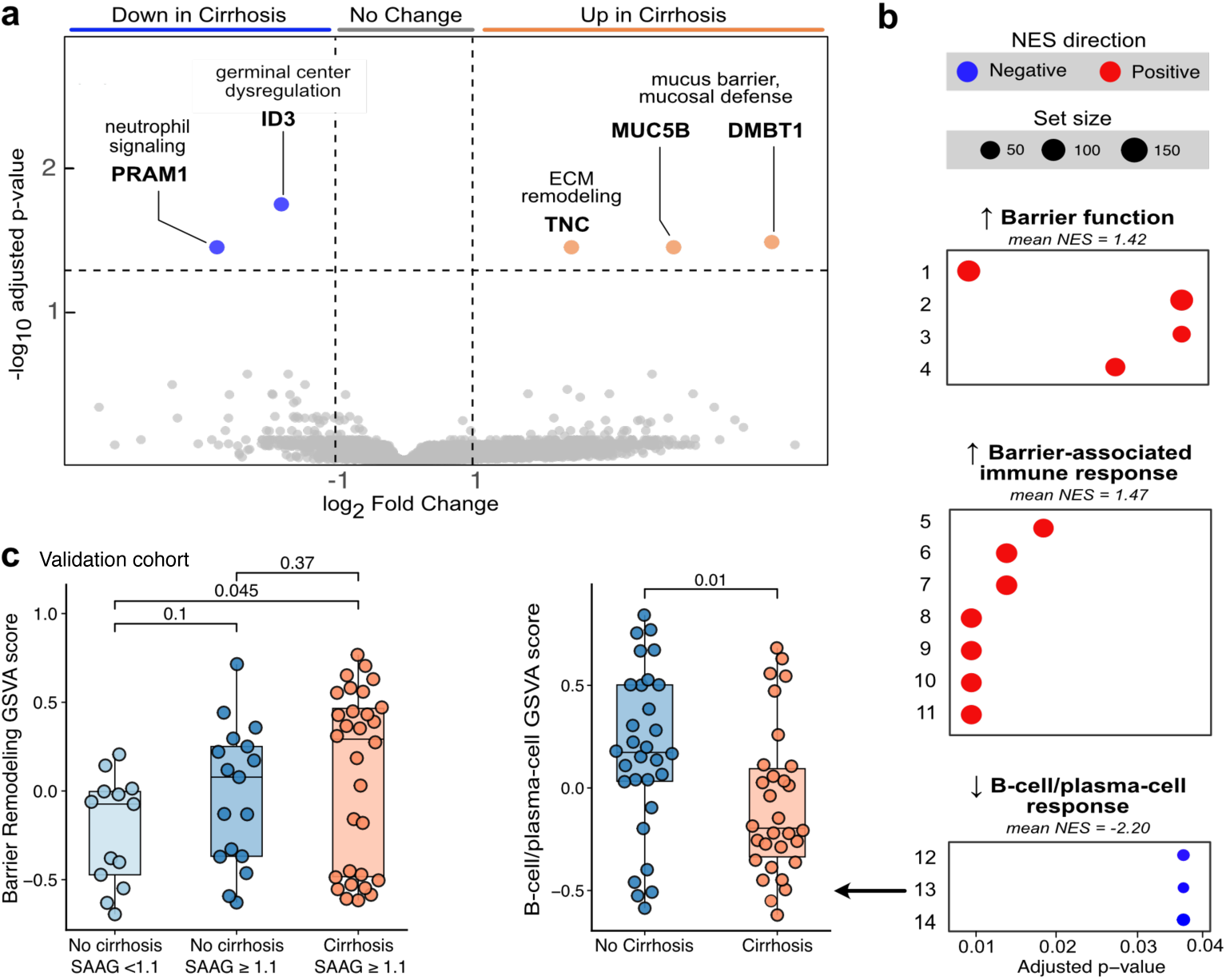
Cirrhotic ascites is characterized by barrier-remodeling and reduced plasma-cell transcriptional programs. (a) Differential host-gene expression in cirrhotic versus non-cirrhotic ascites. Genes with FDR-adjusted P < 0.05 and |log2 fold change| > 1 are highlighted (DESeq2 Wald test with Benjamini-Hochberg correction). (b) Gene set enrichment analysis of the ranked transcriptome. Significant MSigDB gene sets (FDR-adjusted p < 0.05) were grouped into study-defined biological themes. Points are plotted by FDR-adjusted p-value on a shared x-axis across all three themes; color indicates the direction of enrichment, and point size indicates gene-set size. Mean normalized enrichment score (NES) is shown above each theme. (c) Left: validation-cohort GSVA score for the barrier-remodeling program across non-cirrhotic ascites with SAAG <1.1 g/dL (*n* = 13), non-cirrhotic ascites with SAAG ≥1.1 g/dL (*n* = 17), and cirrhotic ascites with SAAG ≥ 1.1 g/dL (n = 30; two-sided Wilcoxon rank-sum test). Right: validation-cohort plasma-cell GSVA score in cirrhotic and non-cirrhotic ascites (two-sided Wilcoxon rank-sum test).

**Figure 3.**
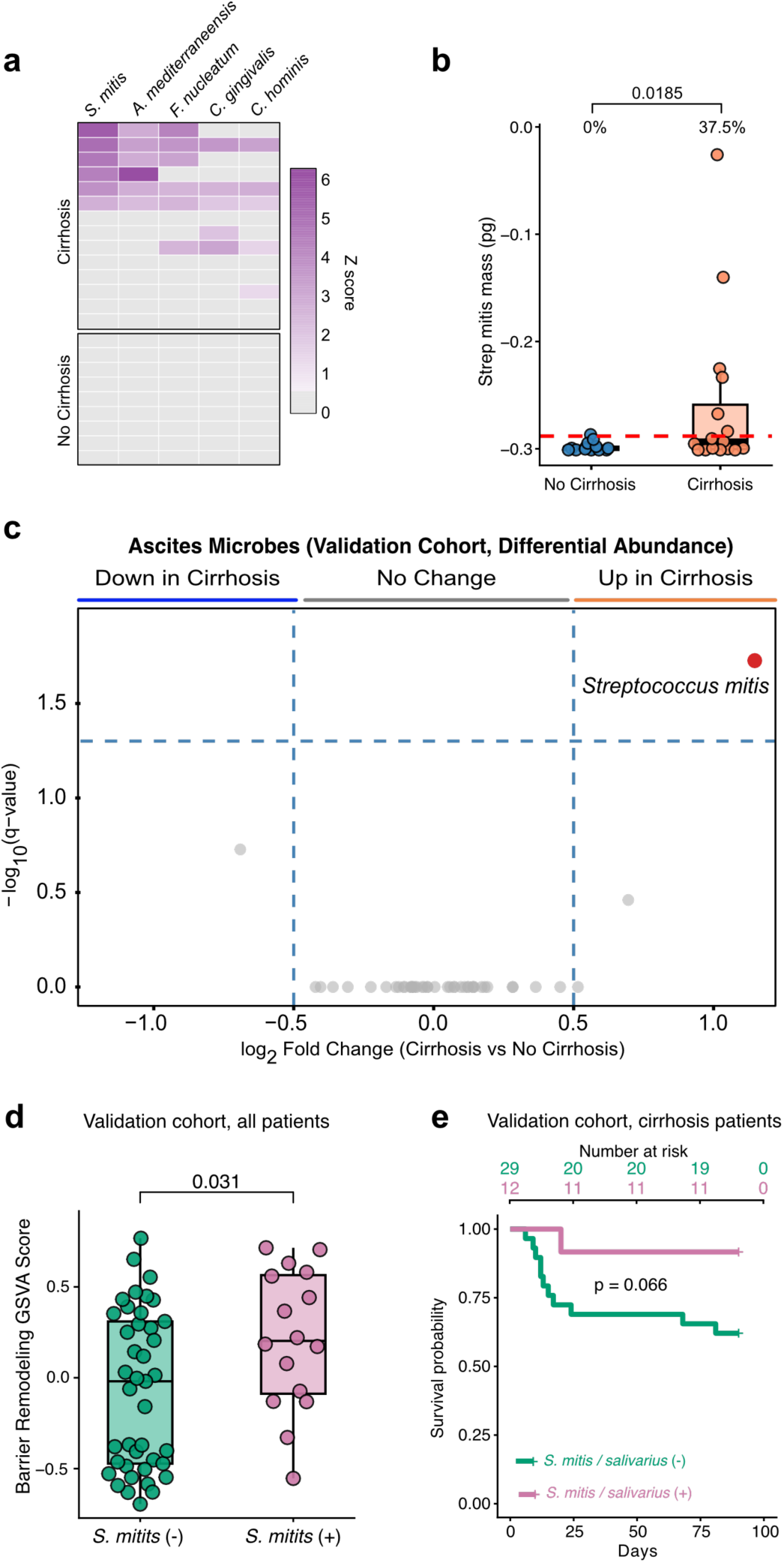
Oral-associated microbes accompany barrier remodeling but do not identify adverse outcomes. (a) Discovery-cohort microbial enrichment Z scores relative to non-cirrhotic ascites showing taxa unique to cirrhotic ascites. (b) Absolute *Streptococcus mitis* mass in discovery-cohort ascites from patients with and without cirrhosis (two-sided Wilcoxon rank-sum test); red line indicates the limit of assay detection. (c) Species-level differential abundance in the validation cohort, with *S. mitis* enriched in cirrhotic ascites; x-axis, log2 fold change; y-axis, FDR-adjusted p-value. (d) Barrier-remodeling GSVA scores according to detectable versus undetectable *S. mitis* (two-sided Wilcoxon rank-sum test) in the validation cohort. (e) Kaplan-Meier estimate of 90-day transplant-free survival among patients with cirrhosis according to detection of *S. mitis or Streptococcus salivarius;* numbers at risk are shown (validation cohort, log-rank test p = 0.066).

**Figure 4.**
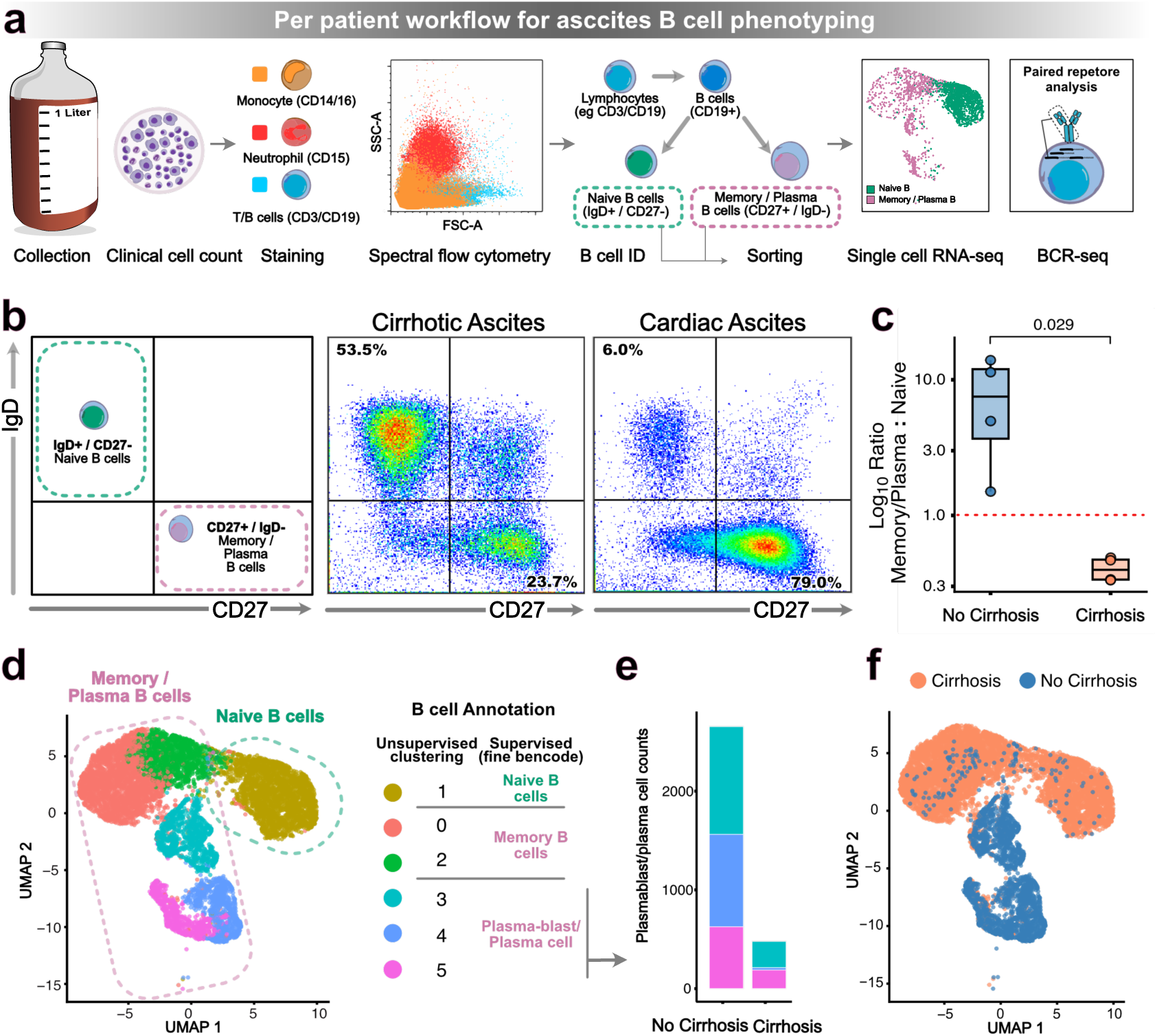
Spectral flow cytometry and single-cell profiling identify remodeling of the ascites B-cell compartment. (a) Workflow for ascites-cell isolation, spectral flow cytometry (*n* = 8), B-cell sorting, single-cell RNA sequencing, and paired BCR sequencing (*n* = 4). (b) Representative CD27 and IgD staining among CD19+ B cells from cirrhotic and non-cirrhotic ascites. (c) Ratio of CD27+IgD-memory/plasma B cells to CD27-IgD+ naive B cells in cirrhotic and non-cirrhotic ascites (*n* = 8); the dashed line indicates a ratio of 1 (two-sided Wilcoxon rank-sum test). (d) UMAP of 18,429 sorted B cells from four patients, colored by unsupervised cluster, with cell-state annotations. (e) Plasmablast/plasma-cell counts in non-cirrhotic and cirrhotic ascites. (f) UMAP colored by cirrhosis status.

**Figure 5.**
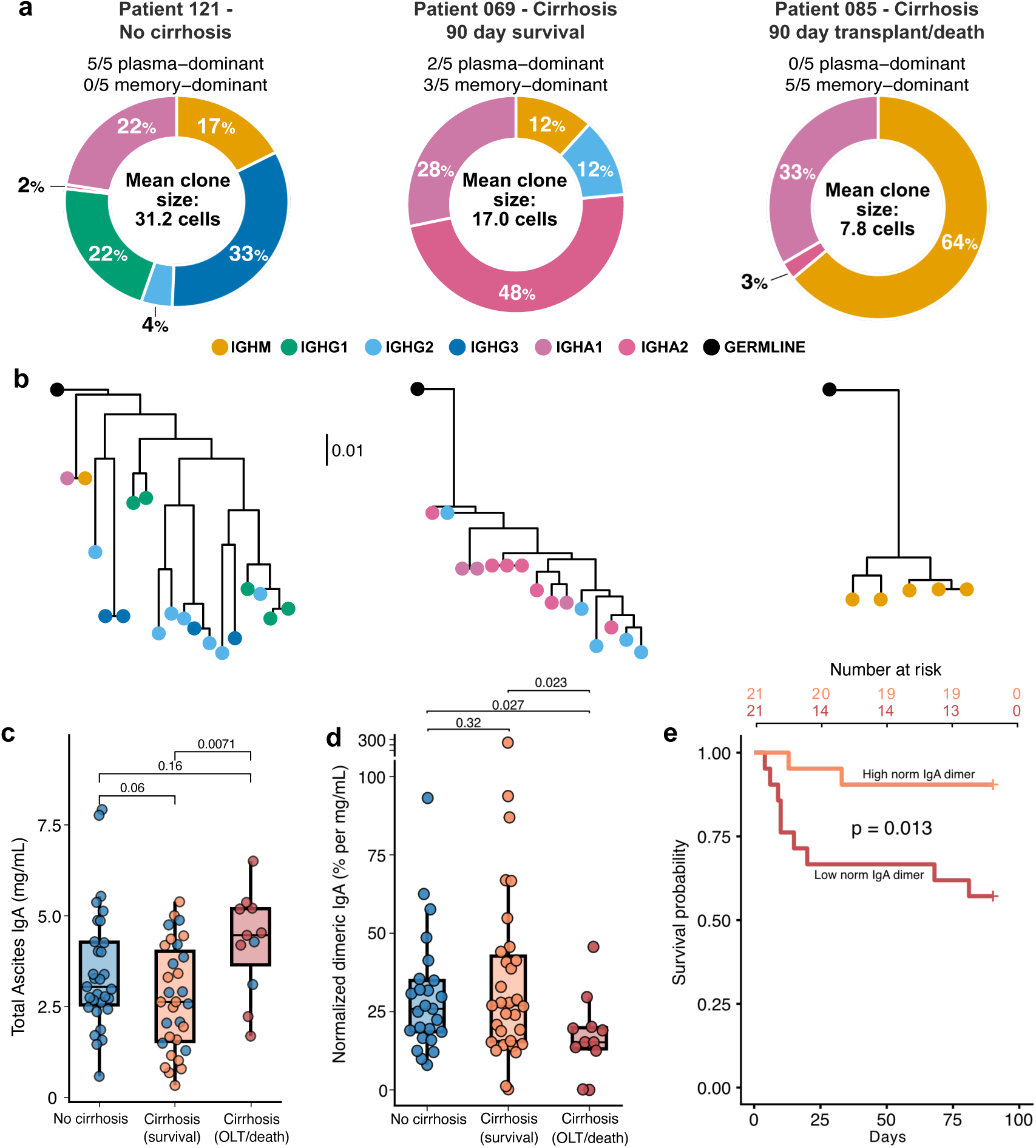
BCR clonal remodeling and reduced dimeric IgA characterize high-risk cirrhotic ascites. (a) The five most abundant BCR clonotypes in three representative patients: no cirrhosis, cirrhosis with 90-day transplant-free survival, and cirrhosis with an adverse 90-day outcome. Donut segments indicate heavy-chain isotype; mean clone size is shown in the donut center, and plasma-dominant (≥50% plasma cells) or memory-dominant clones (≥ 50% memory B cells) are summarized above. (b) Representative BCR lineage trees from dominant clonotypes, illustrating clonal relationships and isotype diversification. (c) Total ascites IgA concentration (two-sided Wilcoxon rank-sum test). (d) Dimeric IgA normalized to total IgA (two-sided Wilcoxon rank-sum test) in the validation cohort. (e) Kaplan-Meier estimate of 90-day transplant-free survival among patients with cirrhosis without spontaneous bacterial peritonitis, stratified by normalized dimeric IgA below versus above the cohort median; numbers at risk are shown (validation cohort, log-rank p = 0.013).

**Table 1a.**
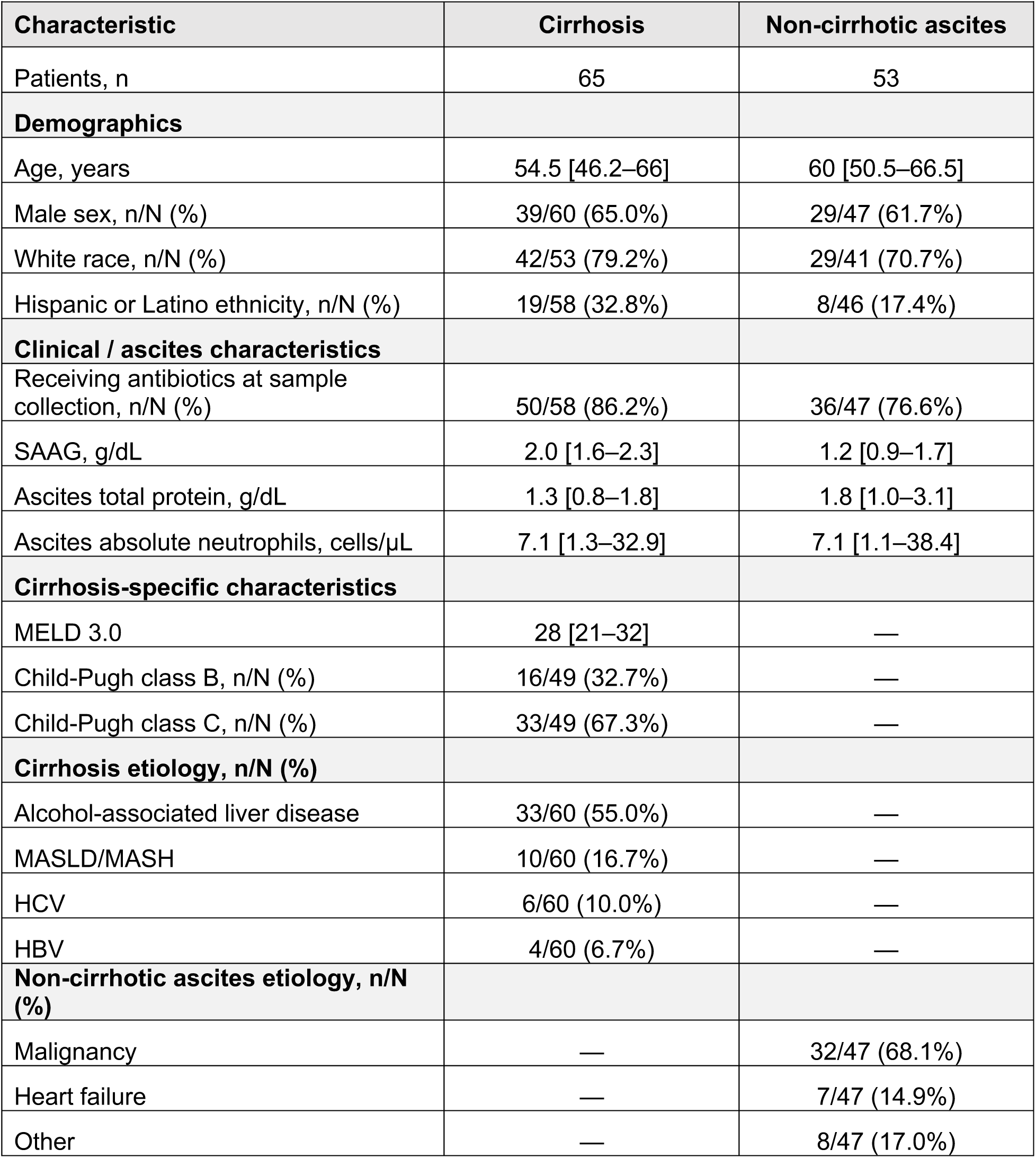
Clinical characteristics of the full study cohort.

**Table 1b.** Clinical characteristics of cirrhosis patients the validation cohort by 90-day transplant-free survival.

| **Characteristic** | **90-day transplant-free survival (n=34)** | **Death or liver transplantation ≤90 days (n=15)** |
| --- | --- | --- |
| Patients, n | 34 | 15 |
| **Demographics** |  |  |
| Age, years | 50 [42–64] | 58 [54–65.5] |
| Male sex, n/N (%) | 18/34 (52.9%) | 14/15 (93.3%) |
| **Cirrhosis severity** |  |  |
| MELD 3.0 | 27.5 [21.5–30] | 30 [22–34] |
| Child-Pugh class C, n/N (%) | 19/34 (55.9%) | 14/15 (93.3%) |
| **Ascites / clinical characteristics** |  |  |
| SAAG, g/dL | 2.1 [1.5–2.4] | 2.0 [1.7–2.1] |
| Ascites total protein, g/dL | 1.2 [0.8–1.7] | 1.6 [1.2–1.9] |
| Antibiotics at sample collection, n/N (%) | 30/34 (88.2%) | 13/15 (86.7%) |
| **90-day outcome follow-up** |  |  |
| Transplant-free survival days | 90 | 13 [10–22] |

Differential gene expression analysis in the discovery cohort identified a limited set of significantly altered genes in cirrhotic relative to non-cirrhotic ascites, with increased expression of genes related to mucus/barrier defense and extracellular-matrix remodeling and decreased expression of genes involved in immune-cell signaling (Figure 2a, Supplemental Table 1a). To identify broader coordinated biological programs, we performed gene set enrichment analysis (GSEA) across the ranked transcriptomic profile. GSEA identified 14 significantly enriched or depleted MSigDB gene sets (adjusted p < 0.05; Supplemental Table 1b). Based on shared biological functions among their leading-edge genes, these gene sets were grouped into study-defined biological themes, revealing enrichment of barrier-function and barrier-associated immune-response programs (mean NES 1.42 and 1.47, respectively) and depletion of B-cell/plasma-cell-associated programs (mean NES −2.20; Figure 2b).

We next examined whether the biological programs identified in the discovery cohort were also evident in a validation cohort of 88 patients. To evaluate these programs at the individual-sample level, we constructed study-defined gene modules by the integrating leading-edge genes identified by GSEA with genes previously associated with the corresponding biological processes in published transcriptional signatures (Supplemental Table 1c). Gene set variation analysis (GSVA) was then used to calculate sample-level enrichment scores for these modules in the validation cohort. Patients were classified into three groups according to the presence of portal hypertension and cirrhosis: non-portal hypertensive ascites (SAAG <1.1 g/dL), portal hypertensive non-cirrhotic ascites (SAAG ≥1.1 g/dL without cirrhosis), and cirrhotic ascites with portal hypertension (SAAG ≥1.1 g/dL with cirrhosis). Barrier-remodeling scores increased across these groups, with significantly higher scores in cirrhotic compared with non-portal hypertensive ascites (p = 0.045; Figure 2c), consistent with progressive remodeling of the ascites microenvironment. Plasma-cell-associated scores were also significantly decreased in cirrhotic ascites (p = 0.01; Figure 2c), recapitulating the suppressed B-cell/plasma-cell program identified in the discovery cohort. Thus, across both discovery and validation cohorts, altered B-cell/plasma-cell biology emerged as an unexpected major theme. In this predominantly uninfected cohort, we did not identify a significant macrophage-associated transcriptional signal, despite macrophages being the most prevalent cell type in ascites and having been implicated in the pathogenesis of SBP^24^.

### Oral-associated microbes accompany barrier remodeling but not adverse clinical outcomes

Microbial translocation in cirrhosis is a known driver of SBP^16–18^, and data from both animal models and humans implicates portal hypertension as a driver of both barrier dysfunction and bacterial translocation^25,26^. We therefore investigated whether portal hypertension was associated with the microbial mass we could detect in ascitic fluid, as a proxy for microbial translocation. To quantify microbial mass in ascites, we applied methods validated for use in low microbial mass settings^5,27^, incorporating extensive environmental controls, and confirmed the sensitivity and reproducibility in ascitic fluid (Supplemental Figure 1, a-d).

When comparing patients with and without cirrhosis, there were no significant differences in overall microbial mass between cirrhotic and non-cirrhotic ascites (Supplemental Figure 1e); however, there was a significant increase in microbial diversity seen in portal hypertensive ascites (Supplemental Figure 1f). In the discovery cohort, we identified a unique microbial composition of cirrhotic ascites (Figure 3a). These distinct microbes are all associated with the oral microbiome, and were enriched both relatively and by absolute mass (Figure 3b).

Analysis of microbial content in ascites from the validation cohort also identified *Streptococcus mitis* to be enriched in cirrhotic patients (Figure 3c). Further, detection of *Streptococcus mitis* (*S. mitis*) was associated with significant enrichment of the barrier remodeling signature (Figure 3d); however, detection of either *S. mitis* or *Streptococcus salivarius (S. salivarius)* was not associated with increased mortality or liver transplant among patients with cirrhosis (Figure 3e). In fact, there was a trend toward lower mortality among patients with *S. mitis* or *S. salivarius* detected in their ascites (p =0.066, Figure 3e). These findings suggested that the detection of characteristic microbes alone does not identify patients with adverse outcomes.

### Cirrhosis is associated with profound remodeling of the ascites B cell compartment

Our transcriptomic analysis identified reduced plasma cell abundance as a prominent feature of cirrhotic ascites. To validate these findings, we isolated cells from 1 liter of ascitic fluid from patients with cirrhosis (*n* = 4) and without cirrhosis (*n* = 4). We developed a spectral flow cytometry assay to phenotype the ascites immune compartment (gating strategy, Supplemental Figure 2) and applied single-cell RNA sequencing (scRNA-seq) to determine whether B-cell populations in cirrhotic ascites were phenotypically distinct from those in non-cirrhotic ascites (Figure 4a). There were no differences in clinical characteristics, including ascites cell count, between patients with cirrhosis (*n* = 4) and without cirrhosis (*n* = 4; Supplemental Table 2 a, b). Patients with cirrhosis demonstrated a significant reduction in the ratio of memory/plasma B cells (CD27+IgD−) to naïve B cells (CD27−IgD+; p=0.029, Figure 4b, c).

We subsequently sorted these phenotypically defined naïve and memory/plasma B-cell populations from four patients (two with cirrhosis and two without) for deep single-cell transcriptomic profiling (Figure 4a). After quality-control filtering, 18,429 B cells were retained for analysis, with comparable cellular yield from cirrhotic and non-cirrhotic patients. Unsupervised clustering identified naïve B cells (cluster 1), two transcriptionally distinct populations of memory B cells (clusters 0 and 2), and three plasmablast/plasma-cell populations (clusters 3–5; Figure 4d); the top marker genes defining each cluster are provided in Supplemental Table 6. Given the depletion of plasma-cell-associated transcriptional programs identified by cohort-wide bulk transcriptomics (Figures 2b, c), we focused subsequent analyses on the plasmablast/plasma-cell compartment.

Cirrhotic ascites contained roughly 10-fold fewer plasmablasts and plasma cells despite comparable overall B-cell sampling between diagnostic groups (Figure 4e). This difference was reflected in a striking population-level shift across the transcriptionally profiled B-cell compartment, with marked loss of the plasmablast/plasma-cell states that prominently characterized non-cirrhotic ascites (Figure 4f). Cluster 5 was a distinct B cell population characterized by high JCHAIN and IGHA1 gene expression, and low MKI67 expression, consistent with a relatively non-proliferative, mucosal-associated plasma-cell phenotype (Supplemental figure 3). As J-chain expression supports assembly of dimeric IgA, the loss of this population suggested that remodeling of the ascites B-cell compartment might be accompanied by altered local IgA responses.

### B-cell remodeling and reduced dimeric IgA characterize high-risk cirrhotic ascites

To more deeply interrogate remodeling of the B-cell compartment in cirrhosis, we evaluated B cell receptor (BCR) sequences and reconstructed lineage trees for the five most abundant clonotypes from each patient (*n* = 4; Supplemental Figure 4, Supplemental Table 3). Within the top 5 expanded clones per patient, we examined clonal abundance, isotype diversification, and B cell subset identification. We summarize these results in three representative patients to illustrate progressive contraction of clonal expansion observed in a patient without cirrhosis, a patient with cirrhosis who remained transplant-free at 90 days, and a patient with cirrhosis who underwent liver transplantation 6 days after study enrollment (Figure 5a). In the two patients with cirrhosis, we observed a reduced number of cells per clone, reduced isotype diversification, and an increasing proportion of clones dominated by IgA and IgM memory responses (Figure 5a). BCR lineage trees similarly showed less isotype diversification among the dominant clonotypes in the cirrhotic patients, including several expanded IgM-restricted memory lineages in the poor-outcome patient (Figure 5b). Importantly, this contraction of the clonally expanded plasma-cell compartment did not reflect an absence of antigen experience. The overall B-cell repertoire was more diverse in cirrhosis (Supplemental Figure 5a), and somatic hypermutation within the plasmablast/plasma-cell compartment was preserved or increased in cirrhosis (Supplemental Figure 5b). Together, these findings suggest an apparent dichotomy between an increased antigen-experienced B-cell repertoire and the reduced abundance of antibody-secreting cells in the local B-cell compartment.

We hypothesized that loss of clonally expanded class-switched plasma cells in ascites would result in quantitative and qualitative differences in local antibody responses. Ascites IgM concentrations were not increased in cirrhosis, consistent with the enrichment of IgM memory B cells, which do not secrete antibodies (Supplemental Figure 6). Given the depletion of JCHAIN-high, IgA-enriched antibody-secreting cells and the preferential production of dimeric IgA by mucosal-associated IgA plasma cells, we measured both total ascites IgA and quantified dimeric IgA using an automated quantitative Western blot assay (Supplemental Figure 7).

Dimeric IgA was normalized to total IgA to estimate the relative abundance of IgA in dimeric form. Total ascites IgA was not reduced in cirrhosis; it was increased among patients who experienced death or liver transplantation within 90 days (Figure 5c); IgG was similarly increased (Supplemental Figure 6). In contrast, normalized dimeric IgA was reduced in patients with adverse outcomes (Figure 5d). Thus, adverse outcomes were not associated with a deficiency of IgA overall, but instead with a shift in the molecular form of IgA present in ascites. Focusing on patients with cirrhosis who did not have SBP (*n* = 42), we found that patients with lower normalized dimeric IgA had worse 90-day transplant-free survival (log-rank p = 0.013; Figure 5e). In an exploratory Cox proportional hazards analysis of these patients, low normalized dimeric IgA remained associated with death or liver transplantation after adjustment for MELD-3.0 (aHR 5.41, 95% CI 1.15–25.37; p = 0.032; Supplemental Table 4).

## Discussion

In this study, unbiased integrated profiling of the ascites microenvironment provided a unified view of host and microbe biology in decompensated cirrhosis, revealing that high-risk disease was distinguished not by greater microbial burden, but by profound remodeling of local humoral immunity. Prior studies have elegantly defined how cirrhosis evolves with increasing disease severity, including changes in peripheral B cell biology^28,29^, and the progressive occupation of the gut microbiome with oral microbes^30–32^. However, disease progression, particularly within decompensated cirrhosis, is both heterogeneous and multifactorial. This inherent complexity poses significant challenges for determining underlying drivers of clinical outcomes. We addressed these challenges by simultaneously interrogating host and microbial features within the shared physiologic context of the ascites microenvironment. This allowed us to resolve interconnected processes and deliver a unified molecular landscape, identifying features of cirrhotic ascites, and linking them to clinical outcomes.

Oral microbes enriched in the gut microbiome of cirrhosis patients have been proposed to contribute directly to decreased barrier integrity and disease progression^32^. Unbiased meta-transcriptomic profiling, which captures both host and microbial changes within the ascites compartment, confirmed a relationship between barrier remodeling and the presence of oral microbes within the ascites. However, the detection of oral-associated microbes in cirrhotic ascites was not associated with adverse outcomes in our study, consistent with prior observations that oral–gut microbial overlap was not associated with 1-year mortality^31^. Notably, although bacterial translocation and infection have long provided the rationale for primary antibiotic prophylaxis against SBP, contemporary evidence^33^ and clinical trials^34^ have not consistently demonstrated a mortality benefit. These findings highlight the need to better understand the factors that shape outcomes among patients with ascites.

Our cohort-wide analysis identified decreased plasma-cell-associated programs as a major feature of cirrhotic ascites compared with non-cirrhotic ascites across diverse etiologies and a range of portal-hypertensive states. This observation was comprehensively validated in focused patient subsets using increasingly high-resolution approaches, including spectral flow cytometry, single-cell transcriptomics, and B-cell receptor sequencing. Prior studies have demonstrated a marked loss of circulating CD27+ memory B cells in cirrhosis.^29,35^. Consistent with these systemic observations, we identified a reduced ratio of CD27+IgD− cells to CD27-IgD+ naïve B cells within cirrhotic ascites. However, higher-resolution profiling revealed that antigen-experienced memory B cells remained well represented in ascites, whereas the most striking alteration was a nearly 10-fold depletion of plasmablasts and plasma cells. Cirrhotic patients maintained a diverse, somatically hypermutated memory B-cell receptor repertoire, suggesting that loss of antibody-secreting cells did not simply reflect an absence of antigen exposure or recognition, but may suggest an impaired local humoral effector response. This distinction may be particularly relevant because differentiated plasma cells preferentially home to and persist within specialized tissue niches, including mucosal plasma cells. ^36,37^.

To evaluate the humoral effector response in ascites, we measured immunoglobulin concentrations and IgA dimerization status. Despite marked depletion of antibody-secreting cells, total ascites IgA and IgG were increased in patients with adverse outcomes, while dimeric IgA was reduced relative to total IgA. Elevated peripheral IgA has similarly been associated with advanced disease and adverse outcomes in cirrhosis^38,39^, raising the possibility that increased circulating immunoglobulin contributes to the ascites antibody pool in high-risk disease.

Because circulating IgA has been reported to be 80-99% monomeric^40,41^, a greater peripheral contribution could expand the total IgA pool while reducing the relative representation of locally produced multimeric IgA. Dimeric IgA is specialized for mucosal transport and immune exclusion at barrier surfaces, making its relative depletion particularly notable in ascites, a compartment positioned at the interface of the gut–liver axis and exposed to translocated microbial products. Our findings suggest that greater immunoglobulin abundance does not necessarily reflect preservation of the local humoral response.

Together, these findings provide convergent evidence across transcriptional, cellular, clonal, and soluble antibody compartments that profound remodeling of the local humoral response is a defining feature of cirrhotic ascites and high-risk disease. Given these findings, we propose a revised model (Figure 6), in which B-cell biology represents a central and underrecognized component of ascites that may underpin poor outcomes in cirrhosis. Non cirrhotic ascites is dominated by a robust population of clonally expanded and class switched plasma cells. In cirrhotic ascites, signatures of barrier remodeling increase with portal hypertension, accompanied by enrichment of oral-associated microbes (Figure 6, middle panel). High-risk cirrhotic ascites is characterized by a divergence between preserved antigen-experienced memory B cells and depletion of class-switched plasma cells, accompanied by a relative decrease in dimeric IgA (Figure 6, right panel). Taken together, these results suggest that adverse outcomes are not determined by microbial exposure alone, but by variation in the local humoral response. Further investigation into the mechanisms underlying this apparent disconnect between antigen experience and effective antibody production may have broader implications for understanding impaired vaccine responsiveness and increased susceptibility to infection among patients with cirrhosis^14,42^.

**Figure 6.**
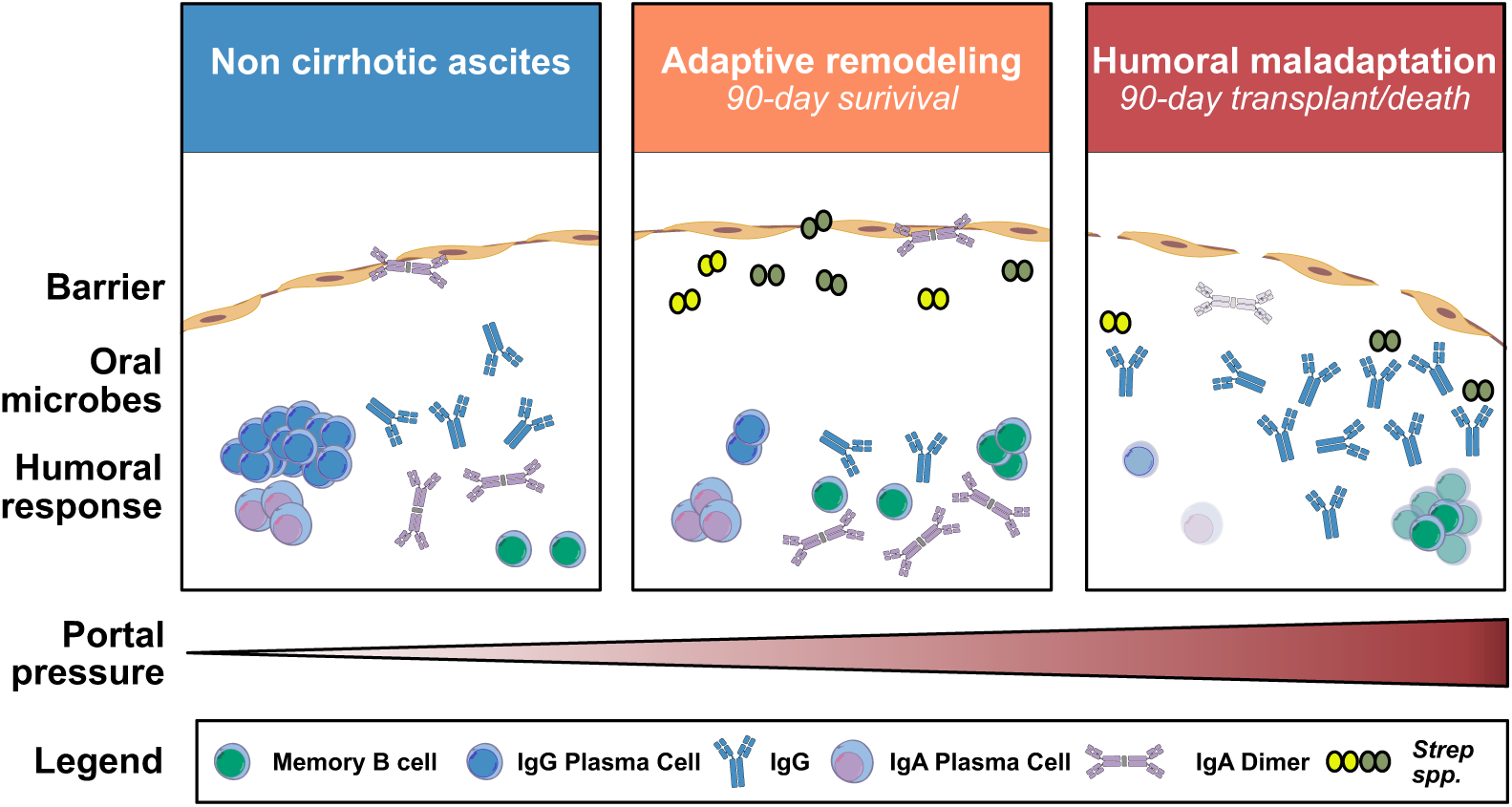
Model of ascites microenvironment remodeling across clinical states. Non-cirrhotic ascites contains clonally expanded, class-switched plasma cells. Cirrhotic ascites is characterized by barrier remodeling, enrichment of oral-associated microbes, and depletion of plasma-cell programs. High-risk cirrhotic ascites shows a divergence between preserved antigen-experienced memory B cells and depletion of class-switched plasma cells, accompanied by reduced dimeric IgA.

This study has limitations. First, all patient samples were derived from a single-center cohort, and the generalizability of these findings requires external validation. Second, although B-cell profiling was performed at considerable depth, orthogonal validation studies using single-cell and B-cell receptor analyses were conducted in much smaller, focused patient subsets (*n* = 4 for single-cell RNA sequencing). Third, microbial detection in ascites is inherently challenging because of its low microbial biomass, compounded by frequent antibiotic exposure, which may have reduced detectable microbial signal. We mitigated these challenges using established approaches for low-biomass microbial sequencing^27^, complementary analytical methods^43^, and experimental validation of assay sensitivity using ultra-low microbial inputs (Supplemental Figure 1). Finally, the limited number of outcome events necessitates caution in interpreting the exploratory Cox proportional hazards analysis, and these outcome associations warrant validation in larger, independent cohorts.

In summary, unbiased profiling of the ascites microenvironment revealed a marked disruption of local humoral immunity in cirrhosis, characterized by depletion of antibody-secreting cells and reduced representation of dimeric IgA, despite evidence of microbial exposure and B-cell antigen experience. These findings suggest that microbial exposure alone may be insufficient to distinguish clinical trajectories in decompensated cirrhosis; rather, differences in the host response to that exposure may be equally important. Whether this humoral phenotype can serve as a prognostic biomarker would require validation in independent cohorts. More fundamentally, determining whether impaired plasma-cell differentiation and decreased dimeric IgA are consequences of advanced disease or contribute directly to its progression may establish whether the local antibody response represents not only a marker of high-risk disease, but a potentially modifiable component of cirrhosis-associated immune dysfunction.

## Methods

### Study design, participants, and clinical outcomes

The Ascites Characterization and Evaluation Study (ACES) is a prospective observational study of adults with ascites undergoing clinically indicated diagnostic or therapeutic paracentesis at the University of California, San Francisco (UCSF). Hospitalized patients scheduled for paracentesis through the UCSF Hospitalist Procedure Service^44^ were prospectively screened for enrollment. Eligibility criteria included age ≥18 years, planned diagnostic or therapeutic paracentesis as part of routine clinical care, and ability to provide informed consent. Individuals who were unable or unwilling to provide informed consent were excluded.

The analytic cohort included 118 participants, including 65 with cirrhosis and 53 without cirrhosis. Analyses were performed in three phases. The discovery cohort included 30 participants, from whom 10 mL of ascites was ultracentrifuged at 50,000 × g for 10 minutes. The validation cohort included 88 separate, subsequently enrolled participants from the same ACES cohort and used 1 mL of neat ascites. A nested deep-phenotyping cohort included eight participants from whom viable cells were isolated from 1 L of fresh ascites; single-cell RNA and paired B-cell receptor sequencing were performed in four participants from this nested cohort.

Clinical information was prospectively collected from the UCSF electronic medical record and linked to study specimens. The primary outcome for survival analyses was death or liver transplantation within 90 days of study enrollment. Analyses of 90-day transplant-free survival were restricted to participants with cirrhosis without SBP, defined as an ascites absolute neutrophil count <250 cells/µL.

The study was approved by the UCSF Institutional Review Board, and all participants provided written informed consent before enrollment using an IRB-approved consent form. Ascites used for research consisted of excess fluid obtained during clinically indicated paracentesis that would otherwise have been discarded; no additional paracentesis was performed for research purposes.

### Ascites processing and RNA extraction

Ascites fluid was collected by sterile technique using a closed, vacuum-sealed collection system and processed in a biosafety cabinet. For the discovery cohort, ascites was aliquoted into sterile ultracentrifuge tubes and centrifuged at 50,000 × g for 10 minutes. Supernatant was removed and retained, and the resulting pellet was resuspended in 1 mL DNA/RNA Shield (Zymo Research). Total RNA was extracted using Zymo Quick-RNA Mini kits according to the manufacturer’s instructions and subsequently used for metatranscriptomic library preparation and sequencing.

### Metatranscriptomic sequencing, taxonomic profiling, and microbial quantification

Metatranscriptomic libraries were prepared using the Watchmaker RNA Library Prep Kit or NEBNext Ultra II RNA library prep kit, according to previously established methods for low-biomass clinical specimens^27,44,45^. ERCC RNA spike-in controls of known input mass were included to permit absolute microbial mass quantification. Meta-transcriptomic sequencing data were processed using the Chan Zuckerberg ID (CZ ID) metagenomic analysis platform^46^. Host and microbial reads were identified using the CZ ID metagenomic pipeline, and resulting taxonomic assignments and nucleotide count metrics were analyzed using custom R workflows adapted from previously described open-source computational pipelines^5^. Taxonomic reports were merged across samples, and genus-and species-level assignments were retained after exclusion of phage assignments, invalid taxonomic identifiers, empty taxonomic categories, and taxa failing prespecified thresholds for nucleotide contig count, alignment length, or nucleotide percent identity. Genus-and species-level nucleotide counts and reads-per-million tables were generated.

According to prior methods^27^, absolute taxon mass was estimated using ERCC spike-ins as: taxon mass = ERCC input mass × taxon nucleotide count / total ERCC reads. Environmental negative controls (water only input, processing controls of just RNAshield, or patient skin swabs from the sterilized paracentesis site post procedure) were used to account for background microbial signal in these low-biomass specimens. For each taxon, the mean and standard deviation of mass measured in negative controls were calculated, and a contamination threshold equal to the control mean plus three standard deviations was subtracted from the measured mass in patient samples; negative adjusted values were set to zero. For microbial enrichment analyses, taxon abundance was log10-transformed after addition of a pseudocount of 1. Sample-level enrichment Z scores were calculated relative to environmental controls as (log10 abundance − control mean)/control standard deviation. Control standard deviations of zero were replaced with 0.0001, and non-finite or missing Z scores were set to zero. Species-level microbial composition in the validation cohort was analyzed using the phyloseq R package, and differential abundance was summarized as log2 fold change with false-discovery-rate-adjusted P values.

### Host gene-expression analysis

Host gene counts were analyzed in R using DESeq2. In the discovery cohort, STAR gene-level count tables were imported using unstranded counts, Ensembl identifiers were mapped to HGNC gene symbols, and duplicate symbols were collapsed by retaining the entry with the greatest total count. Samples with fewer than 10 detected genes were excluded. Genes were retained when at least three samples had ≥10 counts. Prespecified erythrocyte/heme, mitochondrial, and selected housekeeping genes were removed before analysis. In the validation cohort, lowly expressed genes were filtered using edgeR::filterByExpr, and the same filtering was applied. Integer count matrices were modeled with cirrhosis status as the design variable. Variance-stabilizing transformation (VST; blind = FALSE) was used for normalized expression analyses. Differentially expressed genes were defined for visualization as Benjamini-Hochberg FDR-adjusted P < 0.05 and absolute log2 fold change >1.

### Gene-set enrichment and sample-level pathway scoring

Ranked host-gene expression results were analyzed by gene set enrichment analysis (GSEA) using clusterProfiler/fgsea-compatible workflows and MSigDB gene sets. Multiple-testing correction was performed using the Benjamini-Hochberg method, with gene sets meeting an FDR-adjusted p < 0.05 considered significant. Related significant gene sets were grouped into study-defined biological themes based on gene-set context and the leading-edge genes driving enrichment. For validation, study-defined gene modules were constructed by integrating findings from discovery-cohort differential-expression and GSEA/leading-edge analyses with genes associated with the corresponding biological processes in previously described transcriptional signatures (Supplemental Table 1c). Gene set variation analysis (GSVA) was performed on VST-normalized gene-expression data from the independent validation cohort using the GSVA R package to generate sample-level enrichment scores for a barrier-restitution/remodeling module and a plasma-cell module. GSVA was implemented using gsvaParam with a Gaussian kernel (kcdf = “Gaussian”) and a minimum gene-set size of three. Barrier-remodeling scores were evaluated across non-cirrhotic ascites with SAAG <1.1 g/dL, non-cirrhotic ascites with SAAG ≥1.1 g/dL, and cirrhotic ascites with SAAG ≥1.1 g/dL. Plasma-cell GSVA scores were compared between cirrhotic and non-cirrhotic ascites using a two-sided Wilcoxon rank-sum test.

### Immunoglobulin quantitation and measurement of dimeric IgA

Ascites IgA, IgM, and IgG concentrations were measured using Abcam 90-minute ELISA kits according to the manufacturer’s instructions. Ascites samples were diluted 1:10,000 for IgA, 1:1,000 for IgM, and 1:100,000 for IgG and assayed in duplicate. Immunoglobulin concentrations were determined from assay-specific standard curves generated in Microsoft Excel and corrected for the corresponding sample dilution factor.

Dimeric IgA was measured using the JESS automated Western blot platform under nonreducing conditions. Standard solutions were prepared in 0.1× sample buffer, and ascites samples were diluted 1:5 in 0.1× sample buffer. For sample preparation, 2 µL of diluted sample was combined with 2 µL of 0.1× sample buffer and 1 µL of 5× Fluorescent Master Mix prepared without DTT. Water was substituted for DTT solution to maintain nonreducing conditions. Samples were denatured at 70°C for 10 minutes before analysis. Primary antibody was diluted 1:250 in Antibody Diluent 2. Each run included a molecular-weight ladder and common reference samples, including native, serum, and serum:native mixtures at 1:1, 1:9, and 9:1 ratios, to facilitate identification and comparison of immunoglobulin species across runs. Dimeric IgA abundance was normalized to total ascites IgA to generate the normalized dimeric IgA measure used in downstream analyses.

### Spectral flow cytometry and cell sorting

Fresh ascites cells isolated from the nested deep-phenotyping cohort were analyzed by spectral flow cytometry using a 23-marker panel on a Cytek Aurora. The complete antibody panel, including antibody targets and fluorophores, is provided in Supplemental Table 5, and the gating strategy is shown in Supplemental Figure 2. Cells were resuspended in PBS and stained with fluorescent viability dye (FVS780; 1:25,000), followed by washing with FACS buffer to remove excess viability dye. Cells were then resuspended in fresh FACS buffer containing Fc receptor block and stained with the antibody cocktail. Following staining, cells were washed with FACS buffer before acquisition or sorting. The panel was designed to characterize lymphoid and myeloid populations, including T cells, B cells, natural killer cells, monocytes/macrophages, dendritic cells, granulocytes, and plasma cells. CD19+ B cells were evaluated using CD27 and IgD expression, including CD27+IgD− memory/plasma and CD27−IgD+ naïve B-cell populations.

For fluorescence-activated cell sorting, cells were resuspended at approximately 5–9 × 10^6 cells/mL in sorting buffer and passed through a 40-µm cell filter. Cells were maintained on ice and filtered a second time immediately before loading onto the sorter. The sorter was maintained at 4°C with sample mixing enabled, and cells were sorted using Multiway Purity mode into polypropylene collection tubes containing collection buffer. Samples were initially run at flow rate 1, with the flow rate increased as needed while maintaining sorting efficiency above 80%. Flow rate was reduced, or samples were diluted when necessary, to minimize conflict aborts and maintain sorting efficiency. Phenotypically defined B-cell populations were collected for downstream single-cell analysis. An aliquot of sorted cells was analyzed after sorting to assess purity, with the sample line flushed between specimens to minimize carryover.

### Single-cell RNA and paired BCR sequencing

Sorted B-cell populations from four patients (two with cirrhosis and two without cirrhosis) underwent single-cell 5′ gene-expression and paired B-cell receptor (BCR) sequencing using the 10x Genomics Chromium GEM-X 5′ v3 platform. Following fluorescence-activated cell sorting, cells were resuspended in 1× PBS containing 0.04% BSA and concentrated to approximately 1,000 cells/µL for single-cell library preparation. Up to 37.5 µL of cell suspension was used for GEM generation. Gene-expression and BCR libraries were generated from each sample using the Chromium GEM-X 5′ v3 workflow according to the manufacturer’s protocol. Sequencing was performed using the Illumina NovaSeq X.

After sequencing and quality-control filtering, 18,429 B cells were retained for analysis. Single-cell transcriptomic data were analyzed in Seurat using previously established approaches based on the UCSF Wilson Laboratory computational immunology workflow (https://github.com/UCSF-Wilson-Lab/CompImmunologyWorkshopUCSF). Unsupervised clustering and UMAP visualization were performed, and clusters were annotated using canonical B-cell transcriptional programs. Cluster-marker genes were identified using Seurat::FindMarkers with only positive markers, a minimum detection fraction of 0.25, and a log2 fold-change threshold of 0.25.

### B-cell receptor clonotype and lineage analysis

Paired B-cell receptor (BCR) sequencing data were analyzed using previously established computational approaches based on the UCSF Wilson Laboratory computational immunology workflow (https://github.com/UCSF-Wilson-Lab/CompImmunologyWorkshopUCSF), incorporating the Immcantation framework and associated R packages, including alakazam, shazam, dowser, and airr. Briefly, BCR sequences were linked to single-cell transcriptomic metadata by unique cell barcode, and heavy-chain analyses were restricted to productive IGH sequences with assigned clonotypes and available sequence and germline alignments.

Clonal abundance was calculated separately within each participant, and the five most abundant clonotypes from each participant were selected for detailed characterization. Clones were summarized according to cell number, heavy-chain isotype composition, transcriptionally defined B-cell state, and isotype diversification. Somatic hypermutation frequency was quantified using shazam::observedMutations. For phylogenetic analysis, clonotype data were formatted using dowser::formatClones, requiring a minimum of two sequences per clone, and BCR lineage trees were reconstructed using dowser::getTrees and visualized using plotTrees and ggtree. V-and J-gene allele suffixes were removed for visualization and summary of gene usage.

### Clinical and statistical analysis

Continuous variables were summarized as median (interquartile range) and categorical variables as count (percentage). Statistical tests are specified with the corresponding analyses and figure legends when confirmed from the supplied code. Differential-expression and gene-set analyses used false-discovery-rate correction as described above. Ninety-day transplant-free survival was estimated by Kaplan-Meier analysis and compared using the log-rank test.

Cox proportional-hazards models were used for exploratory adjusted survival analyses, including adjustment for MELD-3.0 where specified. All tests were two-sided; P < 0.05 was considered statistically significant unless an FDR-adjusted threshold was specified.

## Data availability

Data underlying the analyses and figures in this study, including processed data and R data objects required to reproduce the reported analyses, will be deposited in the Dryad Digital Repository and made publicly available upon publication. Raw sequencing data will be deposited in appropriate public sequence repositories, with accession numbers provided upon publication. Participant-level clinical data will be shared only in de-identified form and to the extent permitted by participant consent, institutional review board requirements, and applicable protections for participant privacy. Repository DOIs and accession numbers will be provided in the final published article.

## Code availability

Computational analyses in this study were built upon and adapted from previously developed open-source workflows. Single-cell RNA-sequencing and B-cell receptor repertoire analyses were adapted from the UCSF Wilson Laboratory Computational Immunology workflow (https://github.com/UCSF-Wilson-Lab/CompImmunologyWorkshopUCSF). Metatranscriptomic and microbial analyses were adapted from previously established workflows described by Zinter et al.5,27 Study-specific modifications and custom analyses were implemented in R, including workflows for host gene-expression, gene-set enrichment and GSVA, microbial quantification and taxonomic analyses, single-cell and B-cell receptor analyses, and clinical and survival analyses. The R scripts and R Markdown documents containing these study-specific adaptations and analyses, together with associated R data objects required to reproduce the reported results, will be archived in the Dryad Digital Repository upon publication. The Dryad DOI will be provided in the final published article.

