## Supplemental Figures for "Humoral Immune Dysregulation Defines High-Risk Cirrhotic Ascites"

**Supplementary Figure 1. Validation of quantitative meta-transcriptomic sequencing and characterization of microbial burden in ascites.** **a)** Experimental design for validation of microbial detection and quantification in low-biomass ascites. Microbial controls were added at fixed concentrations (1.5–27 fg), and Huh-7.5 cells were added to generate samples spanning medium, low and ultra-low microbial-to-host biomass ratios. **b)** Meta-transcriptomic sequencing and analysis workflow. Samples were sequenced to approximately 50 million reads; host reads were removed and non-host reads were taxonomically annotated. **c)** Measured mass of microbial spike-in controls across medium, low and ultra-low microbial-biomass conditions, demonstrating reproducible quantification across the tested range (intraclass correlation coefficient [ICC] = 0.98). **d)** Reproducibility of microbial community composition across technical replicates from four ascites samples, assessed by Bray–Curtis dissimilarity (ICC = 0.75). **e)** Total microbial mass in ascites according to portal-hypertensive and cirrhosis status. Total microbial mass did not differ significantly among non-cirrhotic ascites with SAAG <1.1 g/dL, non-cirrhotic ascites with SAAG ≥1.1 g/dL, and cirrhotic ascites with SAAG ≥1.1 g/dL. **f)** Shannon diversity according to portal-hypertensive and cirrhosis status. Shannon diversity was higher in cirrhotic ascites with portal hypertension than in non-cirrhotic ascites without portal hypertension (P = 0.01); the comparison between the two non-cirrhotic groups was P = 0.071, and the comparison between portal-hypertensive non-cirrhotic and cirrhotic ascites was not significant. Box plots show the median (center line), interquartile range (box), and 1.5× interquartile range (whiskers); points represent individual participants. Statistical comparisons were performed using two-sided unpaired t-tests.

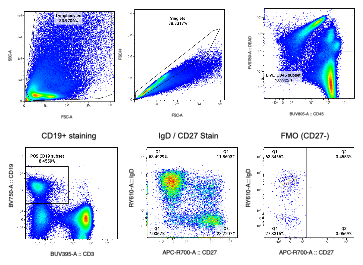

**Supplementary Figure 2. Spectral flow cytometry gating strategy for identification of ascites B-cell subsets.** Representative gating shows sequential identification of total cells, singlets, live CD45+ leukocytes and CD19+ B cells, followed by separation of CD27+IgD− memory/plasma B cells and CD27−IgD+ naive B cells. The CD19+ gate was applied before CD27 and IgD analysis. A representative fluorescence-minus-one (FMO) control for CD27 is shown to illustrate placement of the CD27-positive gate.

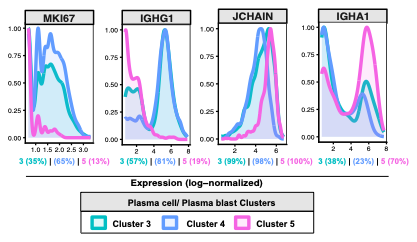

**Supplementary Figure 3. Transcriptional characterization of plasmablast and plasma cell populations.** Log-normalized expression of selected marker genes across clusters 3, 4 and 5 in the four patients who underwent single-cell RNA sequencing. Cluster 3 is shown in green, cluster 4 in blue and cluster 5 in pink. For each gene, expression is displayed among cells positive for the corresponding transcript within each cluster, with the percentage of transcript-positive cells indicated for each cluster. Cluster 5 showed high JCHAIN and IGHA1 expression and low MKI67 expression, consistent with a relatively non-proliferative, mucosal-associated plasma-cell phenotype.

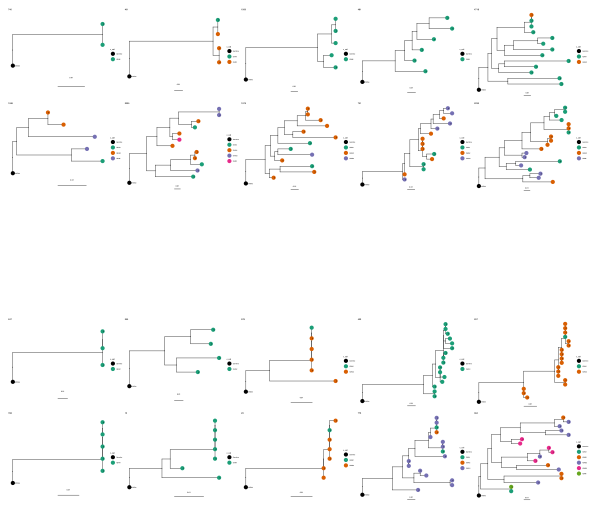

**Supplementary Figure 4. B-cell receptor clonotype and lineage analysis.** BCR lineage trees for the five most abundant clonotypes from each of four deeply profiled patients. Colors denote heavy-chain isotype and may differ among trees. Tree tips are annotated with the transcriptionally defined B-cell cluster of the corresponding B cell when available. Clonotypes are shown separately by participant.

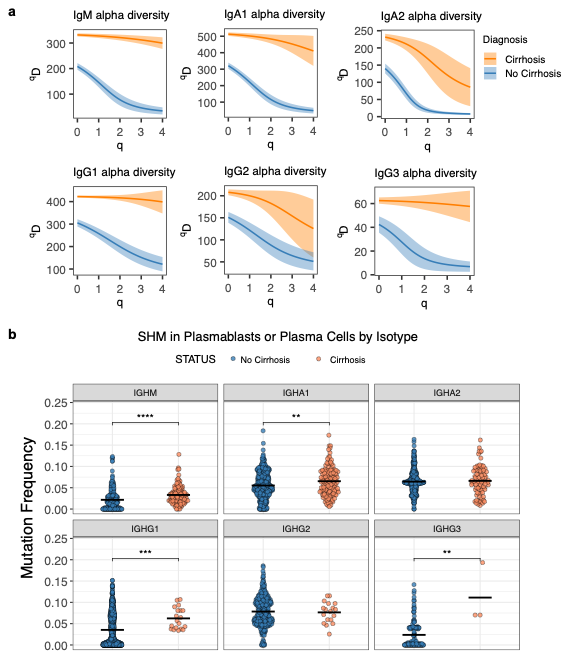

**Supplementary Figure 5. B-cell receptor repertoire diversity and somatic hypermutation.** **a)** Alpha diversity of B-cell receptor repertoires from all analyzed B cells in four patients, shown by BCR isotype and compared between participants with and without cirrhosis. **b)** Somatic hypermutation frequency among cells annotated as plasmablasts or plasma cells, shown by BCR isotype. Statistical significance is indicated by **p < 0.01 and ***p < 0.005.

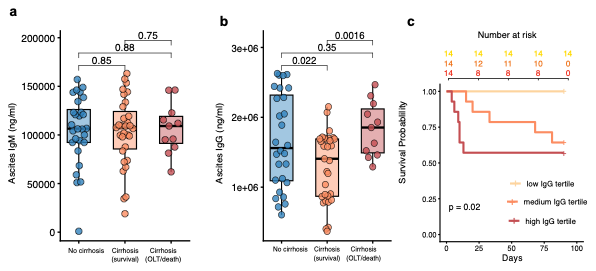

**Supplementary Figure 6. Ascites immunoglobulin concentrations and outcome-associated IgG.** **a)** Ascites IgM concentrations (ng/mL) in participants without cirrhosis, participants with cirrhosis who remained free of death or liver transplantation through 90 days, and participants with cirrhosis who died or underwent liver transplantation within 90 days. **b)** Ascites IgG concentrations (ng/mL) across the same outcome groups. For a and b box plots show the median (center line), interquartile range (box), and 1.5× interquartile range (whiskers); points represent individual participants. Statistical comparisons for a and b were performed using two-sided unpaired t-tests; p values are shown. **c)** Ninety-day transplant-free survival among 42 participants with cirrhosis without spontaneous bacterial peritonitis, stratified by ascites IgG tertile (low, middle and high; n = 14 per group). Survival was estimated by Kaplan–Meier analysis and compared using the log-rank test (P = 0.02).

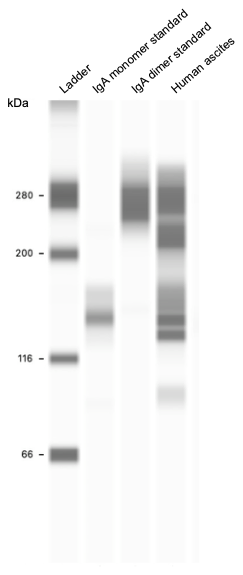

**Supplementary Figure 7. Identification and quantification of dimeric IgA in ascites by automated capillary Western blot.** Representative JESS output showing an IgA monomer standard, an IgA dimer standard and representative ascites samples. IgA species were identified by alignment of sample signal with the molecular-weight positions of the reference standards, and monomeric and dimeric fractions were quantified from the corresponding molecular-weight regions. Dimeric IgA abundance was normalized to total ascites IgA to generate the normalized dimeric IgA measure used in downstream analyses.
